# The septate junction component Bark beetle is required for *Drosophila* intestinal barrier function and homeostasis

**DOI:** 10.1101/2022.11.07.515432

**Authors:** R. A. Hodge, M. Ghannam, E. Edmond, F. de la Torre, Cecilia D’Alterio, N.H. Kaya, M. Resnik-Docampo, T. Reiff, D. L. Jones

## Abstract

Age-related loss of intestinal barrier function has been found across species, and the causes remain unknown. The intestinal epithelial barrier is maintained by tight junctions (TJs) in mammals and septate junctions (SJs) in insects. Specialized tricellular junctions (TCJs) are found at the nexus of three adjacent cell membranes, and we showed previously that aging results in mis-localization of the tricellular SJ (tSJ) component Gliotactin (Gli) in enterocytes (ECs) of the *Drosophila melanogaster* intestine. In embryonic epithelia, the tSJ protein Bark beetle (Bark) recruits Gli to tSJs, which prompted us to investigate Bark function in the intestine. Bark protein localization decreases at tSJs in aged flies. EC-specific *bark* depletion in young flies led to hallmarks of intestinal aging and shortened lifespan, whereas depletion of *bark* in progenitor cells reduced Notch activity, biasing differentiation toward the secretory lineage. Together, our data implicate Bark in EC maturation, maintenance of intestinal barrier integrity, and homeostasis. Understanding the assembly and maintenance of tSJs to ensure barrier integrity may lead to strategies to improve tissue integrity when function is compromised.

## Introduction

The intestinal barrier allows selective paracellular transport of water, ions, and nutrients while maintaining food and microbes inside the intestinal lumen (Marchiando *et al*., 2010). Barrier integrity is crucial to intestinal homeostasis because leakage of potentially harmful antigens or microbes could be detrimental to interstitial tissues. There is a strong correlation between aging and decline in intestinal barrier integrity across multiple species, including *Drosophila melanogaster* (Kirkwood, 2004; Biteau *et al*., 2008; Ren *et al*., 2014; Schiffrin *et al*., 2010; Tran & Greenwood-Van Meerveld, 2013; Rera *et al*., 2012). Age-associated loss of the intestinal barrier in *Drosophila* is associated with systemic metabolic defects, such as changes in insulin/insulin-like growth factor signaling, intestinal dysbiosis, chronic expression of inflammatory genes, and an increase in proliferation of intestinal stem cells (ISCs) (Clark *et al*., 2015; Rera *et al*., 2012; Guo *et al*., 2014). Intercellular occluding junctions, referred to as tight junctions (TJs) in vertebrates and septate junctions (SJs) in arthropods, play a major role in maintenance of the intestinal barrier (Resnik-Docampo *et al*., 2017; Salazar *et al*., 2018; Izumi *et al*., 2019). While TJs and SJs appear different ultrastructurally, they share many homologous proteins, and both TJs and SJs restrict passive, paracellular transport. TJs use a branched network of independently sealing strands to create a semipermeable barrier at “kissing points” between adjacent membranes, while SJs reduce net solute diffusion by increasing diffusional distance across the junction with bridge-like structures called septa (Mariano *et al*., 2011; Furuse & Tsukita, 2006). In *Drosophila*, pleated SJs (pSJs) are found in epithelia derived from ectoderm, whereas smooth SJs (sSJs) are found in endoderm-derived tissues, such as the midgut (Tepass & Hartenstein, 1994; Lane & Skaer, 1980; Resnik-Docampo *et al*., 2018). Similar to TJs, sSJs in *Drosophila* are located apical to adherens junctions (Resnik-Docampo *et al*., 2018), making the *Drosophila* midgut a tractable model system for studying mammalian TJs, such as those that maintain the mammalian intestinal barrier.

The *Drosophila* posterior midgut (PMG) is functionally analogous to the human small intestine (Apidianakis & Rahme, 2011). The PMG epithelium is composed primarily of absorptive enterocytes (ECs), polyploid cells with microvilli that extend into the gut lumen and facilitate the absorption of nutrients. ECs and secretory enteroendocrine (EE) cells are maintained by a pool of ISCs that are able to self-renew by producing new ISCs. ISCs primarily generate daughter enteroblasts (EBs), which then differentiate into either mature ECs or EE cells (Apidianakis & Rahme, 2011). EB fate is specified by activation of the Notch signaling pathway (Ohlstein & Spradling, 2007; Micchelli & Perrimon, 2006), leading to the expression of the transcription factor *klumpfuss (klu)* in EBs. EE cell and EC fate decisions become stochastic upon loss of lineage specifying *klu*, and EB fates can be tracked using specific *klu*-based lineage tracing tools (e.g. *klu*^*ReDDM*^) (Reiff *et al*., 2019; Korzelius *et al*., 2019).

In aged flies, ISC proliferation increases dramatically, as measured by an increase in cells that are positive for the mitotic marker phospho-histone H3 (pHH3). (Biteau *et al*., 2008; Li & Jasper, 2016; Jiang *et al*., 2009; YJ *et al*., 2008; Park *et al*., 2009). The increase in progenitor cells is coupled with a delay in differentiation, resulting in an accumulation of cells co-expressing the ISC/EB markers Escargot, Delta, and reporters of the Notch signaling pathway (Biteau *et al*., 2008). Smooth SJs are found between adjacent ECs and between ECs and EE cells, and our lab has demonstrated previously that integrity of sSJs between ECs is required for maintenance of the intestinal barrier in the midgut (Resnik-Docampo *et al*., 2017). Thus, additional intestinal phenotypes exhibited by aged flies include loss of sSJ physical integrity (visible gaps between membranes) (Resnik-Docampo *et al*., 2017) and loss of the intestinal barrier (Rera *et al*., 2012). Indeed, electron micrographs of ECs in aged flies revealed gaps between adjacent ECs, suggesting a loss of sSJ integrity over time (Resnik-Docampo *et al*., 2017). Gaps in SJs correlated with changes in localization of key sSJ components in PMGs from old flies. Immunofluorescence (IF) imaging revealed decreased staining intensity for several sSJ components at the junction, relative to the cytoplasm, with a concomitant increase in staining intensity in the cytoplasm, relative to what was observed for cells in intestines from young flies (Resnik-Docampo *et al*., 2017).

In *Drosophila*, specialized junctions, referred to as tricellular junctions, are found where three adjacent cells meet. Similar to sSJ proteins, the tricellular septate junction (tSJ) protein Gliotactin (Gli) was lost from the tSJ and increasingly localized to cytoplasmic puncta in ECs from aged flies (Resnik-Docampo *et al*., 2017). Bark beetle (Bark), another tSJ-specific protein, also referred to as Anakonda, is required for proper Gli localization to the tSJ in the fly embryonic epithelium. Bark and Gli colocalize at the tSJ in both pSJs and sSJs (Byri *et al*., 2015; Hildebrandt *et al*., 2015). Because Gli is required at tSJs between ECs to maintain intestinal homeostasis (Resnik-Docampo *et al*., 2017), and because proper Gli localization was disrupted in aged flies, we wanted to determine whether Bark also plays a role in the adult intestine. Here, we show that Bark localizes to the tSJ and that it is required in both EBs and mature ECs to maintain intestinal homeostasis.

## Results

### Bark is expressed in the adult intestine

Bark is a type I transmembrane protein with a tripartite extracellular domain; it has been proposed that Bark forms a trimer, which acts as a diaphragm in the center of the tSJ canal (Byri *et al*., 2015). Bark was demonstrated to be required for proper localization of Gli and maturation of the pSJ in the embryonic epithelium (Byri *et al*., 2015). In order to determine whether Bark has a role in the adult intestine, PMGs from young (2 do) flies expressing a GFP-tagged form of Bark (hereafter referred to as Bark::GFP, see Materials and Methods) (Sarov *et al*., 2016) were stained with anti-Bark antibody (Hildebrandt *et al*., 2015). Bark localized to tSJs in both the PMG and hindgut, similar to what was observed in the embryonic epithelium (Byri *et al*., 2015) **(Figs 1, EV1, and 2A)**. Specifically, in the PMG, Bark localized to the tSJ between adjacent ECs and ECs and EE cells. By contrast, in aged (40 day old, do) flies, Bark localization decreased at the tSJ between adjacent ECs and appeared to spread along the bicellular junction, with modest accumulation in the cytoplasm (**Fig 2B,B’,C**). Loss of Bark from the tSJ in PMGs from aged flies is similar to what was observed for Gli; however, previous studies indicated significantly more accumulation of Gli in the cytoplasm, when compared to Bark (Resnik-Docampo *et al*., 2017).

**Fig 1:**
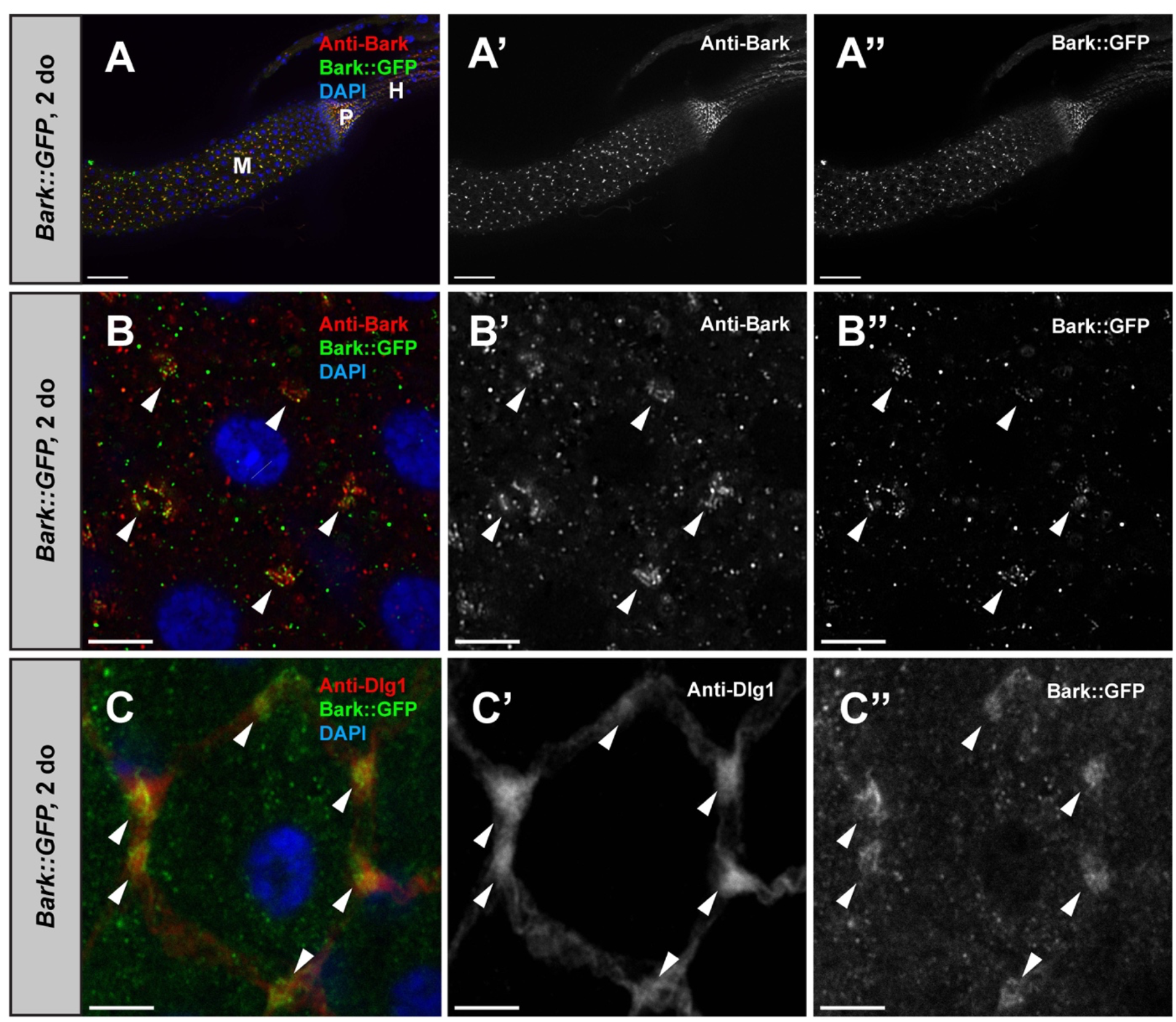
Anakonda/Bark beetle (Bark) localizes to the tricellular septate junction (tSJ) in the adult *Drosophila* intestine. (**A-A”**) Representative staining for Bark (antisera, red) (*Bark::GFP*, GFP, green) in the posterior midgut (M) of young (2 do) flies. Pylorus (P) and hindgut (H) to the right. DNA is stained with DAPI (blue). Scale bars, 50 μm. (**B-B”**) High magnification view of an enterocyte (EC) showing staining of endogenous Bark protein and GFP-tagged bark at the tSJ (arrowheads). (**C-C”**) High magnification view of an EC showing GFP-tagged bark colocalizes with the bicellular septate junction (bSJ) protein Discs large 1 (Dlg1, red) at the tSJ (arrowheads). Scale bars, 5 μm.

**Fig 2:**
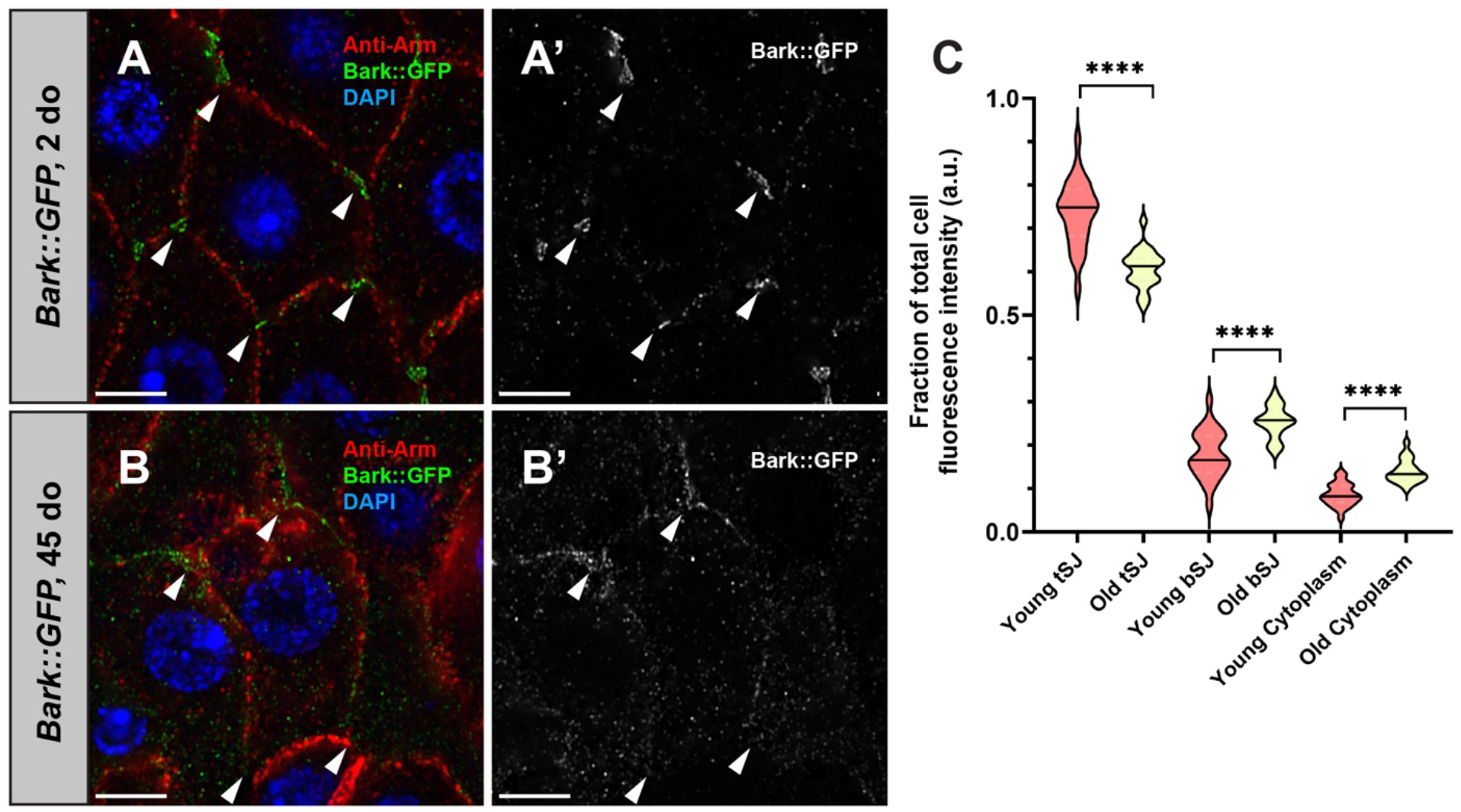
Bark is lost from the tSJ in aged flies. **A)** In young (2 do) flies, Bark localizes to the tSJ (arrowheads) in ECs, whereas it is no longer detected at the tSJ in the aged (45 do) flies **(B-B’)**. Adherens junctions (Armadillo, Arm, red); Bark (GFP, green); DNA (DAPI, blue). Scale bars, 5 μm. (**C**) Quantification reveals that Bark is increased at the bicellular septate junction (bSJ) or in the cytoplasm of old flies. 3 cells were analyzed per PMG. Young *Bark::GFP* PMGs, n = 26; old *Bark::GFP* PMGs, n = 22. 3 cells were measured per PMG. Each clearly visible tSJ in the cell was measured. 6 measurements were taken of the BCJ and cytoplasm in each cell, respectively. Quantification of fluorescence intensity ratios for Bark (**C**) in the bicellular junction (BCJ), tSJ, and cytoplasm. Statistical significance determined by Mann-Whitney test. ns, not significant; *, *P* < .05; **, *P* < .001; ****, *P* < .0001.

Given the previously described relationship between Bark and Gli, where Bark is required for recruitment of Gli to the tSJ in developing embryonic epithelia (Byri *et al*., 2015), we wanted to determine whether Bark is required for Gli localization in the adult PMG. As mutations in *bark* are embryonic lethal, we took advantage of an inducible gene expression system, GeneSwitch (GS), to deplete *bark* from ECs, specifically in young adult flies. The GeneSwitch (GS) system permits cell type-specific gene expression following feeding flies the progesterone analog mifepristone (RU-486) (Brand & Perrimon, 1993; Osterwalder *et al*., 2001). Similar to previously published results in the embryonic epithelium (Byri *et al*., 2015), RNAi-mediated, EC-specific depletion of *bark* resulted in a decrease in Gli::GFP staining intensity at the tSJ (**Fig EV2A-B**). However, in contrast to the findings in the embryonic epithelium, depletion of *Gli* using a similar strategy resulted in an increase in Bark::GFP staining intensity at the bicellular junction, somewhat reminiscent of the aging phenotype (**Fig EV2D-E**). A similar mutually dependent relationship between Gli and Bark has been observed in the pupal notum (Esmangart de Bournonville & le Borgne, 2020). In addition, both, Bark and Gli-depletion, non-autonomously increase ISC proliferation (**Fig EV2C,F**). Taken together, these data suggest that the mechanisms for assembly and maintenance of the tSJ may differ by tissue type or with age.

### bark is required in ECs for the maintenance of intestinal homeostasis

Given that the changes in Gli::GFP upon depletion of *bark*, and vice versa, resemble what occurs at the tSJ in an aged midgut, we wanted to determine whether depletion from ECs in PMGs from young flies led to other aging phenotypes. Following depletion of *bark* from ECs for 5 days in young flies, the number of pHH3^+^ cells was significantly higher than in controls (**Fig 3A-C**), suggesting that Bark is required in ECs to maintain intestinal homeostasis, similar to Gli (Resnik-Docampo *et al*., 2017). Immunostaining confirmed depletion of Bark protein from ECs, validating the RNAi line **(Fig EV3A-B)**. In addition, the number of pHH3^+^ cells increased with extended induction times (**Fig EV3E-G**), and similar results were obtained using additional, independent *RNAi* lines **(Fig 3C, Fig EV3C-D**), as well as alternative, inducible systems to deplete *bark* from ECs **(Fig EV3H-K)**. As a result of increased ISC proliferation, depletion of Bark in EC also increases total cell number **(Fig EV3J)** and progenitor cell numbers shown by esg^+^ cells **(Fig EV3K)**, suggesting disrupted intestinal homeostasis.

**Fig 3:**
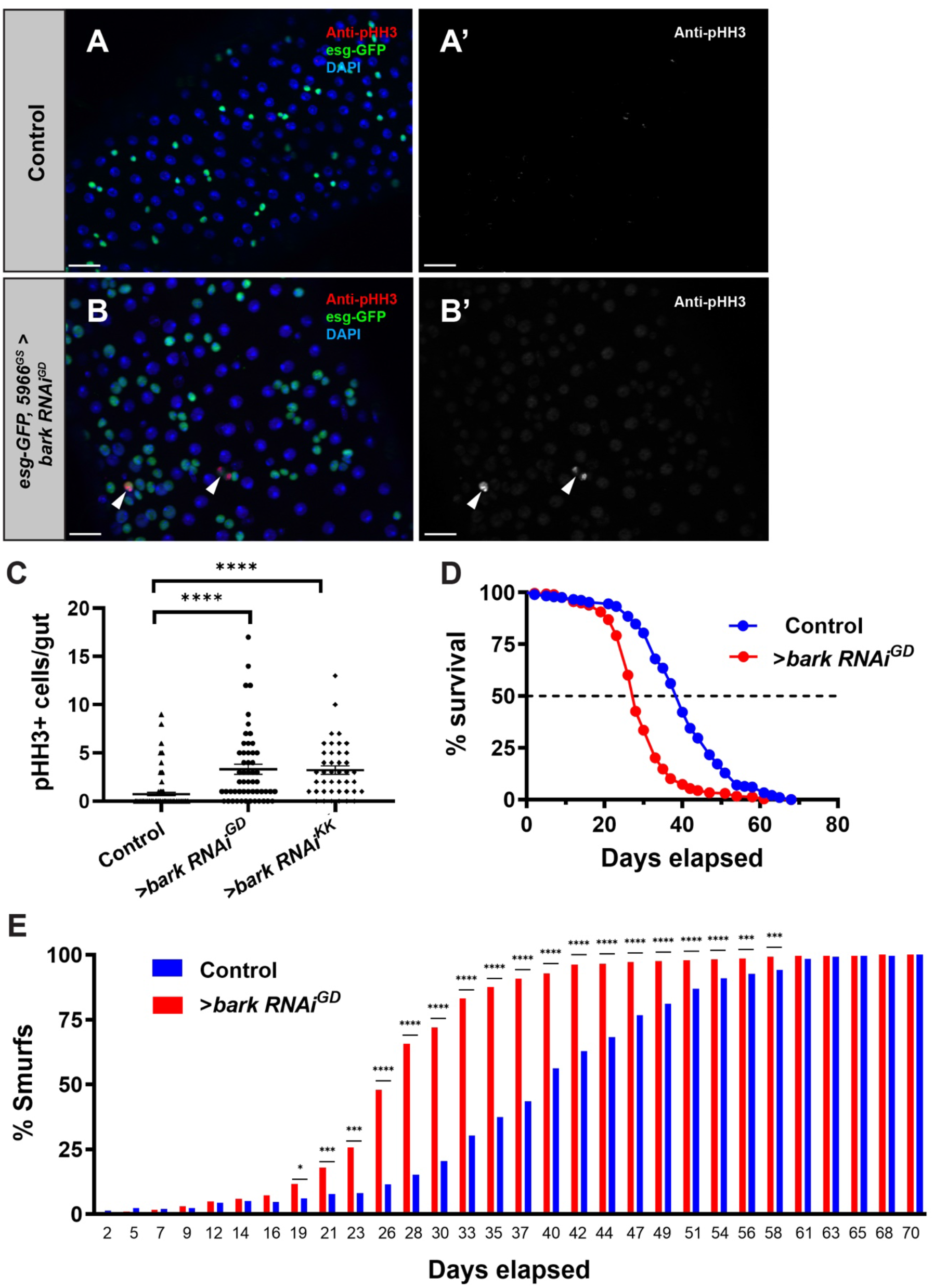
Bark is required at the tSJ in enterocytes to maintain intestinal homeostasis. (**A-B**) Depletion of Bark in ECs of young (2 do) flies by *5966*^*GS*^*:bark RNAi*^*GD*^ expression for 5 days results in an increase in mitotic activity (n = 54), compared to outcrossed controls (n = 48). Genotypes: (**A**) *w; esg-GAL4, UAS-GFP, 5966*^*GS*^. (**B**) *w; esg-GAL4, UAS-GFP, 5966*^*GS*^*:bark RNAi*^*GD*^. All flies are fed RU-486 to induce transgene expression. ISCs/EBs (GFP, green); mitotic cells (phosphorylated histone H3, pHH3, red), DNA (DAPI, blue). Scale bars, 20 μm. (**C)** Quantification of mitotic events. *w; esg-GAL4, UAS-GFP, 5966*^*GS*^*:bark RNAi*^*KK*^ flies are also included (n = 42). Error bars represent mean with SEM. Significance was determined by Mann-Whitney test. ****, *P* < .0001. (**E**) Intestinal barrier integrity assay shows loss of barrier function upon depletion of Bark (red, n = 301) and (**D**) statistically significant (****) shortening of lifespan, when compared to outcrossed controls (blue, n = 299). Genotypes: Red, *w; esg-GAL4, UAS-GFP, 5966*^*GS*^*>UAS-bark RNAi*^*GD*^; blue, *w; esg-GAL4, UAS-GFP, 5966*^*GS*^. *, *P* < .05; ***, *P* < .001, ****, *P* < .0001. Statistical significance determined by Fisher’s exact test (barrier integrity assay) and non-parametric log-rank Mantel-Cox test (lifespan assay).

Next, we wanted to determine whether Bark is required for maintenance of the intestinal barrier, similar to Gli (Resnik-Docampo *et al*., 2017). The GeneSwitch system was used to deplete *bark* in ECs throughout the adult lifespan. Integrity of the intestinal barrier and lifespan were assayed in parallel by feeding the flies a non-absorbable, non-toxic blue food dye, as described previously (Rera *et al*., 2011, 2012). In flies with an intact intestinal barrier, the dye remains in the lumen of the digestive tract; however, when the intestinal barrier is severely compromised, the dye visibly leaks into the hemolymph (Rera *et al*., 2011). Consistent with a model where Bark plays an integral role in tSJs, depletion of *bark* in ECs led to significant loss of the intestinal barrier after 19 days. Furthermore, as has been shown previously, loss of barrier integrity correlated with shortened lifespan when compared to control flies (**Fig 3D-E**) (Rera *et al*., 2012). These data indicate that Bark is required in ECs for regulating intestinal homeostasis, maintaining the intestinal barrier, and for a normal lifespan.

### bark is required in EBs for proper differentiation

A hallmark of the aging PMG is accumulation of EB-like cells that express hallmarks of both ISCs and ECs (Biteau *et al*., 2008; Jiang *et al*., 2009). Although Bark expression was detected at tSJs between ECs and EEs, single cell sequencing data revealed expression of *bark* in EBs (Dutta *et al*., 2015; Hung *et al*., 2021). Therefore, we wanted to investigate a role for *bark* in committed EC progenitors using lineage tracing in combination with conditional expression of *bark* RNAi. Briefly, *klu*^*ReDDM*^ allows EB-specific manipulation and tracing of differentiated ECs, taking advantage of different fluorophore stability **(Fig 4A)** (Reiff *et al*., 2019; Korzelius *et al*., 2019). Depleting *bark* in *klu*^*ReDDM*^ results in accumulation of *klu*^*+*^ EBs (**Fig 4B-D**), although not at the expense of EC differentiation (**Fig 4E**). Similar results were obtained when *bark* was depleted from ISC and EB using *esg*^*ReDDM*^ (**Fig EV4**) (Antonello *et al*., 2015). As a consequence of disrupted cellular homeostasis, specific depletion of *bark* from EBs reduced lifespan compared to outcrossed controls (**Fig 4G**). In addition, depletion of *bark* in EBs led to an increase in newly generated EEs, identified by antibody staining for the EE cell lineage marker Prospero (Pros^+^, **Fig 4F**) and non-autonomously increased ISC proliferation (pHH3^+^, **Fig 4H-J**). Supporting our observations in ageing EC **(Fig 3C)**, EB-specific Bark-depletion reduces immunofluorescence of the SJ marker Dlg1 at bSJ (**Fig 4K-M**).

**Fig 4:**
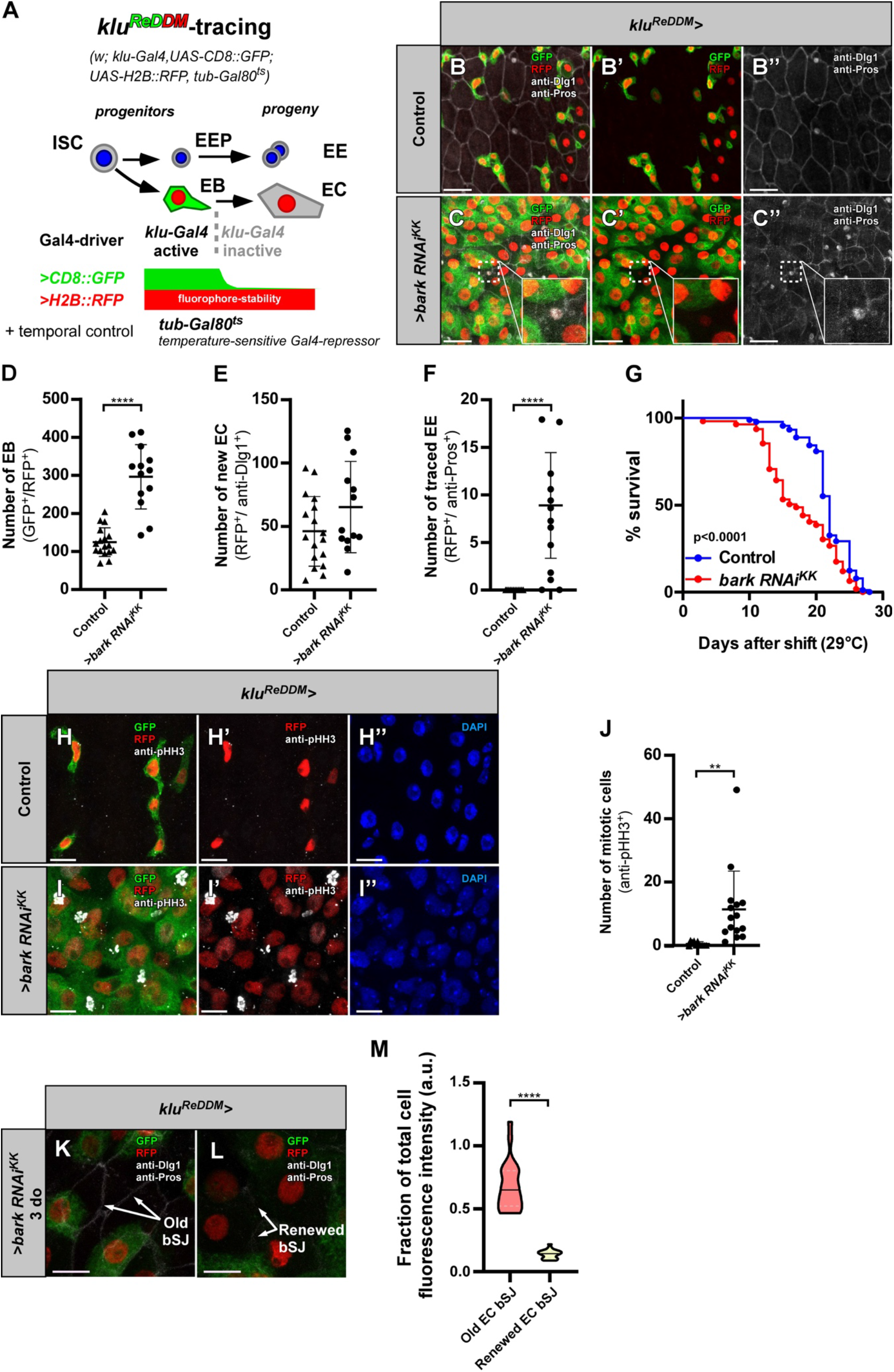
EB specific depletion of bark results in EB accumulation, EE fate plasticity and reduced Dlg1 levels at bSJ of new EC. **(A)** *klu*^*ReDDM*^ (*w; klu-GAL4, UAS-CD8::GFP; UAS-H2B::RFP, tub-GAL80*^*ts*^) tracing. Expression of two fluorophores (*CD8::GFP* and *H2B::RFP*) is driven by EB-specific driver *klu-GAL4*. EBs differentiating to epithelial ECs lose *CD8::GFP*, while stable *H2B::RFP* persists. The expression of UAS-driven transgenes is temporally controlled by a ubiquitously expressed temperature-sensitive *GAL80*^*ts*^ repressor, which is inactivated by temperature shift to 29°C. **(B-C’’)** Confocal images of control PMG **(B-B’’)** and >*bark RNAi*^*KK*^ **(C-C’’)** in the R5 region after 7 days of tracing. **(D-F)** Quantifications of the number of EBs **(D)**, new ECs **(E)** and traced EEs **(F)** upon *bark* knockdown after 7 days of tracing. (control n = 17, *>bark RNAi*^*KK*^ n = 13). Statistical significance determined by Mann-Whitney test. **, *P* < .01. Scale bar 20 μm. Depletion of *bark* induces EB to EE differentiation (inset, GFP-/RFP^+^/anti-Pros^+^, **F**) but no significant change in the number of ECs (GFP^-^/RFP^+^/ anti-Dlg1^+^, **E**). Additionally, *bark* knockdown increased the number of progenitor cells (GFP^+^/RFP^+^, **D**). **(G)** Survival (in percentage) over time upon *bark* knockdown using *klu*^*ReDDM*^. Survival curves were plotted combining data from >80 flies per one genotype group: Control (blue, n = 89), *bark RNAi*^*KK*^ (red, n = 109). *bark* knockdown in EBs reduces lifespan significantly. Statistical significance determined by non-parametric log-rank Mantel-Cox test. ****, *P* < .0001. **(H-I’’)** Knockdown of *bark* **(I-I’’)** leads to an increase in the number of proliferating cells (marked by pHH3) as compared to the controls **(H-H’’). (J)** Quantification of the number of mitotic cells upon *bark* depletion after 7 days of tracing. (control n=11, *>bark RNAi*^*KK*^ n = 15). Statistical significance determined by Mann-Whitney test. ****, *P* < .0001. Scale bar 10 μm. **(K, L)** Knockdown of *bark* leads to a reduced Dlg1 fluorescence intensity at the bSJ between renewed EC (white arrows, **L**) as compared to the Dlg1 fluorescence intensity bSJ between old EC (white arrow, **K**). **(M**) Quantification of the Dlg1 fluorescence intensity ratio at the bSJ between old ECs and bSJ between renewed ECs upon *bark* knockdown after 3 days of tracing. (control n = 24, *>bark RNAi*^*KK*^ n = 22). Statistical significance determined by Mann-Whitney test. ****, *P* < .0001. Scale bar 10 μm.

Cell fate decisions after ISC division strongly depend on the Notch signaling pathway, whereby low Notch activation was shown to drive EE differentiation (Micchelli & Perrimon, 2006; Ohlstein & Spradling, 2007, 2005). By using *Su(H)-GBE-GFP* as a reporter of Notch activation (Furriols & Bray, 2000), we found that depletion of *bark* using *klu-GAL4* led to a reduction in *Su(H)-GBE-GFP* fluorescence intensity (**Fig 5A-C)** and smaller EB nuclei (**Fig 5D)**. Our data indicates that upon *bark* depletion, EBs demonstrate an increase in differentiating along the EE lineage, which has previously been observed upon loss of the lineage marker *klu* in EBs (Reiff *et al*., 2019; Korzelius *et al*., 2019). Taken together, our findings suggest that Bark plays a role during EB to EC differentiation and that depletion of *bark* leads to accumulation of EBs with reduced Notch activity, increasing differentiation towards EE fate.

**Fig 5:**
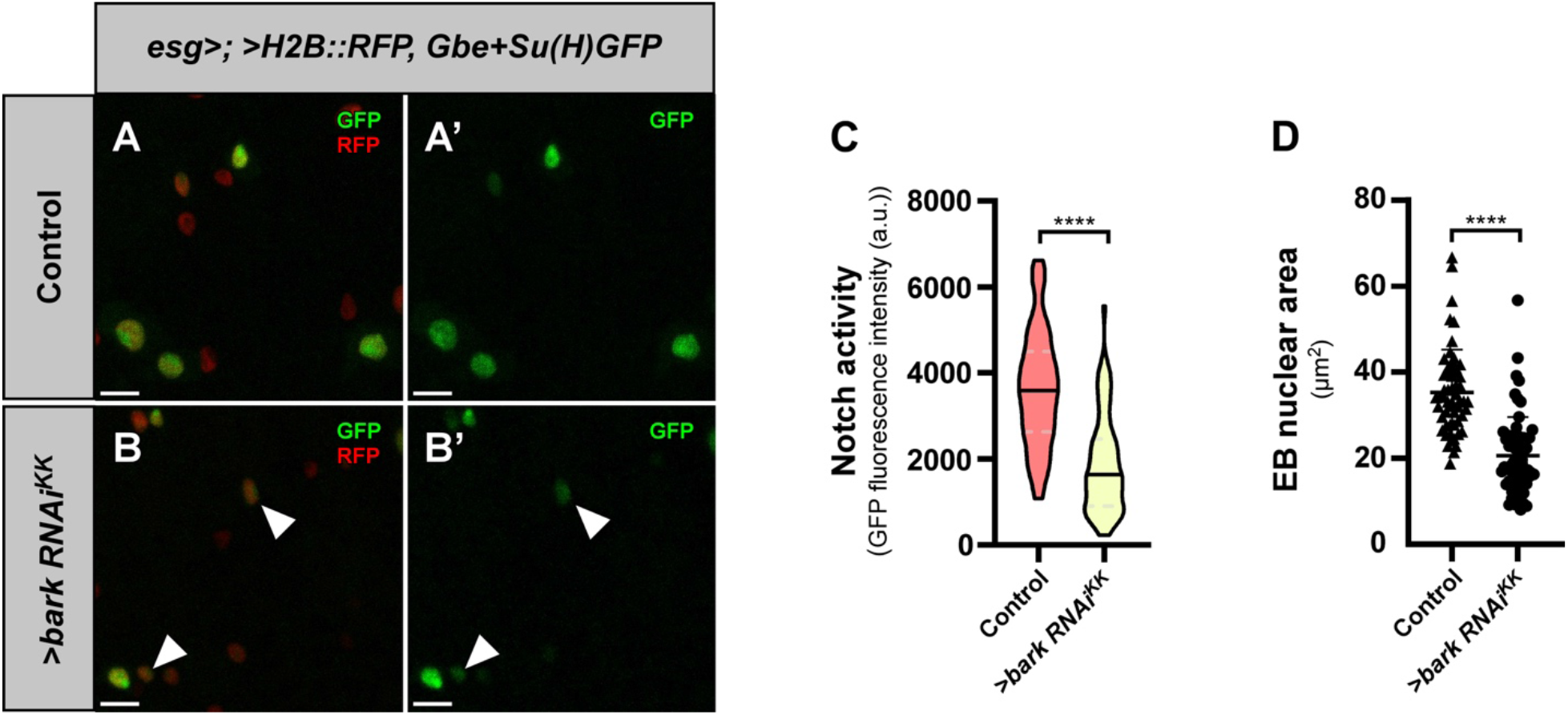
Knockdown of *bark* in ISCs/EBs results in reduced Notch pathway activation and EB nuclear size. **(A)** Expression of *H2B::RFP* fluorophore is driven by ISC- and EB-specific driver *esg-GAL4*. H2B::RFP persists in epithelial ECs differentiated from EBs. *Gbe+Su(H)GFP* is a readout of Notch activity, and marker of the EB lineage (arrowheads, B); GFP intensity was analyzed upon *>bark RNAi*^*KK*^ (**B**). **(C, D)** Quantification of GFP fluorescence intensity and **(D)** EB nuclear size following 7 days of induction. Knockdown of *bark* decreases the GFP intensity (**C**) and EB nuclear size (GFP^+^ cells, **D**) compared to controls (**B, B’**, control n = 73, *>bark RNAi*^*KK*^ n = 70). Statistical significance determined by Mann Whitney test. ****, *P* < .0001. Scale bar 10 μm.

## Discussion

Occluding junctions, known as tight junctions in mammals and septate junctions in arthropods, play a significant role in maintaining the intestinal barrier (Marchiando *et al*., 2010; Resnik-Docampo *et al*., 2017; Salazar *et al*., 2018), which passively restricts paracellular flow across the intestinal epithelium. Interestingly, loss of the intestinal barrier has been described as a result of aging and is associated with impending death across species (Rera *et al*., 2012). Due to the strong correlation between aging and loss of the intestinal barrier (Kirkwood, 2004; Biteau *et al*., 2008; Ren *et al*., 2014; Schiffrin *et al*., 2010; Tran & Greenwood-Van Meerveld, 2013; Rera *et al*., 2012), we wanted to characterize the impact of aging on occluding junctions, using the *Drosophila* intestine as a model system. Here, we describe the expression, localization and role of the tSJ component Bark in the adult PMG.

Bark localized to the tSJ in both the PMG and hindgut (**Figs 1, EV1**); however, analysis of PMGs from aged flies indicated that Bark was lost from the tSJ over time (**Fig 2**). Concomitant with the decreased staining intensity at the tSJ, we observed an increase in staining along the bicellular junction (**Fig 2**). Depletion of *bark* from ECs in young flies led to an increase in ISC proliferation, loss of the intestinal barrier, and shortened lifespan (**Fig 3**). These results are similar to what we reported previously for another tSJ component, Gli (Resnik-Docampo *et al*., 2017). Although our study utilizes RNAi-mediated depletion of tSJ components, we have shown previously that transcription of *bark* does not decrease with age in the PMG (Resnik-Docampo *et al*., 2017); therefore, our approach may not accurately recapitulate the impact of aging on the tSJ. However, these studies are a relevant first step toward understanding the importance of Bark and other SJ proteins in intestinal homeostasis.

Our finding that Bark is lost from the tSJ in PMGs from aged flies, in combination with the observation that depletion of *bark* from ECs in young flies recapitulates some prevalent intestinal phenotypes that are apparent with age, supports the hypothesis that loss of SJ components may contribute to age-related loss of intestinal barrier function (Resnik-Docampo *et al*., 2017; Salazar *et al*., 2018). One possible mechanism leading to age-related changes in the localization of SJ proteins could be alterations in vesicular trafficking, including increased endocytosis of SJ proteins and/or decreased delivery of recycled or newly synthesized SJ proteins to the plasma membrane. Manipulation of transport machinery, rather than production of SJ proteins, could be explored as an approach to restore Bark and other SJ proteins at the SJ in aged flies, although manipulation of transport machinery would likely result in pleiotropic, unrelated phenotypes.

In addition to analyzing a role for Bark in ECs, we show that EB-specific depletion of *bark* results in accumulation of EB-like cells, resembling defects observed in flies that are aged or have damaged intestinal epithelia (Biteau *et al*, 2010; Zhai *et al*, 2015). The accumulation of EB-like cells correlated with a decrease in lifespan (**Fig 4D, G**), as reported previously (Antonello *et al*., 2015). EBs depleted for *bark* are capable of forming ECs in normal numbers (**Fig 4E**), although the new ECs, originating from Bark-depleted EBs, are abnormally small (**Fig 5A-B, D)**, show irregular Dlg1 localization **(Fig 4K-M)** and plasticity is shifted towards EE fate **(Fig 4F)**. Therefore, we speculate that depletion of *bark* from EBs results in failure to form a tight, functional bSJ and/or tSJ, although it was not possible to assess whether normal levels of Bark protein are found in ECs derived from *klu+* EBs/EC progenitors. Our findings demonstrate an essential role for Bark during tSJ establishment when differentiating EBs integrate into the intestinal epithelium. Important future studies will focus on an ultrastructural study of Bark localization in late EBs, achieving a better understanding of how Bark may participate in recently identified apical membrane initiation sites (Chen and St. Johnston, 2022) while EBs undergo epithelial integration, and simultaneously preserve intestinal barrier function, will.

In our evaluation of the relationship between Gli and Bark protein in the PMG tSJ, we found that depletion of one component results in changes in staining intensity of the other at tSJs in adult PMGs (**Fig EV2)**. Our findings are different from previous studies that examined the relationship between Bark and Gli in the embryonic epithelium (Byri *et al*., 2015), which found Bark is required for Gli to localize to the TCJ, whereas the reverse was not true. However, another study revealed Gli and Bark are mutually dependent for localization to the tSJ in the pupal notum (Esmangart de Bournonville & le Borgne, 2020). Differences could be due to relationships between Gli and Bark in embryonic versus adult tissues, the use of null alleles versus RNAi-mediated depletion, length of RNAi depletion time, and/or the use of antibodies targeting the proteins directly, rather than GFP tags. An ultrastructural study could better pinpoint the nature of Gli-Bark interaction in the PMG. Investigation of Gli and Bark’s relationship to another tSJ protein, M6 (Wittek *et al*, 2020; Esmangart de Bournonville & le Borgne, 2020) in the *Drosophila* PMG may also help further illuminate these differing results.

In summary, we have found that Bark protein is required in the tSJs of ECs and EBs in the *Drosophila* posterior midgut to maintain homeostasis in the contexts of ISC proliferation, EB fate, intestinal barrier function, and lifespan. Because Bark also mislocalizes from the tSJ in the aged fly PMG, our study suggests that this mislocalization may contribute to the overall intestinal aging phenotype. Additionally, Bark and the tSJ protein Gliotactin appear mutually dependent on one another for proper tSJ localization in the adult PMG, unlike in the embryonic epithelium (Byri *et al*., 2015). Given the importance of intestinal barrier function, and age-related intestinal barrier loss across species (Kirkwood, 2004; Biteau *et al*., 2008; Ren *et al*., 2014; Schiffrin *et al*., 2010; Tran & Greenwood-Van Meerveld, 2013; Rera *et al*., 2012), our discovery of an essential role for Bark during homeostatic cellular turnover sheds light on fascinating future research lines. This study could help generate insight into the role of tSJs in the human gut and possible pathologies associated with tSJ dysfunction.

## Materials and Methods

Set 1: Fig 1C, Fig 4-5, Fig EV3H-K, Fig EV4

Set 2: Fig 1A-B, Fig 2-3, Fig EV1-EV2, Fig EV3A-G

### Fly food and husbandry

#### Set 1

Fly food contained 7.12% corn meal, 4.5% malt extract, 4% sugar beet syrup, 1.68% dried yeast, 0.95% soy flour, 0.5% agarose, 0.45% propionic acid, and 0.15% NIPAGIN powder (antimycotic agent). *GAL80*^*ts*^ flies were kept at 18°C (permissive temperature) until shifted to 29°. Otherwise, flies and crosses were kept at 25°C. All analyses were performed on 3-7 day old (do) mated females that were shifted to 29°C for 7 days for tracing. For survival analysis (**Fig 4G**, *klu*^*ReDDM*^ *bark* knockdown), 3-5 do flies (males and females) were shifted to 29°C and monitored for their survival.

#### Set 2

Flies were cultured in vials containing standard cornmeal medium (1.1% agar, 2.9% baker’s yeast, 9.2% maltose, and 7.1% cornmeal; all concentrations given in wt/vol). Propionic acid (0.5%) and TegoSept (methylparaben, Sigma, 0.16%) were added to adjust pH and prevent fungal growth, respectively. Newly eclosed adults were kept for an additional 1–2 days before inducing transcription activation by placement on food containing the steroid hormone mifepristone (RU-486; Sigma M8046) in a 25 μg/ml concentration and flipped every 2 days thereafter. All analyses for these studies were performed on female flies, as age-related gut pathology has been well established in females (Biteau *et al*., 2008; Rera *et al*., 2012).

### Fly stocks used

Fly stocks used:
| Stock | Source |
| --- | --- |
| 5966-GAL4<sup>GS</sup> | Gift from H. Jasper<sup>1</sup> |
| *esg-GFP; UAS-H2B::RFP* | T. Reiff |
| *Gbe<sup>+</sup>Su(H)dsRed* | Gift from T. Klein<sup>2</sup> |
| *Gli::GFP* | DGRC 115-332 |
| *Myo1A-Gal4* | Gift from H. Jasper<sup>1</sup> |
| *P{bark.Ty1-Tev-SGFP-FLAG}* | Gift from S. Luschnig<sup>3</sup> |
| *Su(H)lacZ; esg::GFP, 5966GAL4<sup>GS</sup>* | Gift from B. Ohlstein<sup>4</sup> |
| *tub-GAL80<sup>ts</sup>* | BL 7017 |
| *UAS-bark-RNAi<sup>GD</sup>* | VDRC 52608GD |
| *UAS-bark-RNAi<sup>KK</sup>* | VDRC 107348KK |
| *UAS-Gli RNAi* | VDRC 37115GD |
| *UAS-N-RNAi* | VDRC 14477GD |
| *w; esg-Gal4,UAS-CD8::GFP; UAS-H2B::RFP, tub-Gal80ts* | T. Reiff |
| *w; klu-Gal4,UAS-CD8::GFP; UAS-H2B::RFP, tub-Gal80ts* | T. Reiff |
| *w; klu-Gal4,UAS-CD8::RFP; UAS-H2B::RFP, tub-Gal80ts* | T. Reiff |
| *W<sup>1118</sup>* | Gift from L. Wang<sup>5</sup> |
| *yw hsflp; Sco/Cyo Dfd-GMR nuGFP; UAS-Aka attP2y (L057)* | Gift from S. Luschnig<sup>3</sup> |

<sup>1</sup>Genentech and Buck Institute for Research on Aging, USA; <sup>2</sup>University of Düsseldorf, Germany;
<sup>3</sup>University of Münster, Germany; <sup>4</sup>UT Southwestern Medical Center, USA; <sup>5</sup>UCLA, USA

### Immunohistochemistry

#### Set 1

Dissected guts were fixed in 4% PFA in 1XPBS for 45 min. After fixation the guts were washed with 1XPBS for 10 min and stained with primary antibodies, diluted in 0.5% PBT (0.5% Triton (Sigma-Aldrich) in 1XPBS) + 5% normal goat serum (Thermo Fisher Scientific, Berman, Germany). Primary antibody staining was performed at 4°C overnight on an orbital shaker. On the next day, guts were washed with 1XPBS for 10 min and incubated with secondary antibodies and DAPI (1:1000; 100 μg/mL stock solution in 0.18 M Tris pH 7.4; DAPI No. 18860, Serva, Heidelberg) for at least 3 h at RT. After washing with 1XPBS for 10 min the stained guts were mounted in Fluoromount-G Mounting Medium (Electron Microscopy Sciences). Stained posterior midguts were imaged in the R5 region using LSM 710 confocal microscope (Carl Zeiss) using ‘Plan-Apochromat 20 × /0.8 M27′ and ‘C-Apochromat 40 × /1.20 W Corr M27′ objectives. Image resolution was set to at least 2048 × 2048 pixels. Focal planes were combined into Z-stacks and images were then processed by Fiji software. Final images were assembled using Canvas X-Pro. We would like to acknowledge the Center for Advanced Imaging (CAi) at Heinrich Heine University for providing support with imaging and access to the LSM 710 microscope system (DFG INST 208/539-1 FUGG).

#### Set 2

All images were taken in the P3–P4 regions of the *Drosophila* PMG, located by centering the pyloric ring in a ×40 field of view (fov) and moving 1–2 fov toward the anterior. PMGs were dissected into ice-cold PBS/4% formaldehyde and incubated for 1 h in fixative at room temperature. Samples were then washed three times, for 10 min each, in PBT (PBS containing 0.5% Triton X-100), 10 min in Na-deoxycholate (0.3%) in PBT (PBS with 0.3% Triton X-100), and incubated in block (PBT-0.5% bovine serum albumin) for 30 min at room temperature. Samples were immunostained with primary antibodies overnight at 4ºC, washed 3 × 10 min at room temperature in PBT (PBS containing 0.5% Triton X-100), incubated with secondary antibodies (1:500, Invitrogen) at room temperature for 2 h, washed 3 × 10 min with PBT and mounted in Vecta-Shield/DAPI (Vector Laboratories, H-1200). Images were acquired on a Zeiss LSM780 or LSM880 inverted confocal microscope (UCLA MCDB/BSCRC Microscopy Core), and on a Zeiss Axio Observer Z1 with Apotome 2 using the ZEN Black v.2.0 (Zeiss) software. Images were processed with Fiji/ImageJ and Zen software. Final figures were assembled using Adobe Illustrator.

| Antibody | Manufacturer | Catalog # | Species | Clonality | Dilution |
| --- | --- | --- | --- | --- | --- |
| Anti-Armadillo | DSHB | N2 7A1 | Mouse | Monoclonal | 1:20 |
| Anti-Bark | Gift from R. Schuh<sup>1</sup> | n/a | Guinea pig | Polyclonal | 1:1000 |
| Anti-Dlg1 | DSHB | 4F3 | Mouse | Monoclonal | 1:250 |
| Anti-GFP | Aves Labs | GFP-1010 | Chicken | Polyclonal | 1:500 |
| Anti-GFP | Abcam | ab13970 | Chicken | Polyclonal | 1:200 |
| Anti-GFP | Molecular Probes | A-11122 | Rabbit | Polyclonal | 1:3000 |
| Anti-pHH3 | Millipore | 06-570 | Rabbit | Polyclonal | 1:200 |
| Anti-Pros | DSHB | MR1A | Mouse | Monoclonal | 1:250 |
| Anti-Chicken Alexa488 | Invitrogen | A-11039 | Goat | Polyclonal | 1:500 |
| Anti-Guinea Pig Alexa568 | Invitrogen | A-11075 | Goat | Polyclonal | 1:500 |
| Anti-Mouse Alexa568 | Invitrogen | A-11004 | Goat | Polyclonal | 1:500 |
| Anti-Mouse Alexa647 | Invitrogen | A-21235 | Goat | Polyclonal | 1:500 |
| Anti-Rabbit Alexa647 | Invitrogen | A-31573 | Donkey | Polyclonal | 1:500 |

<sup>1</sup>Max Planck Institute for Multidisciplinary Sciences, Germany

### Statistical Analysis

GraphPad Prism 8.0.2 was used to run statistical analysis and create graphs of quantifications. For single comparisons, data sets were analyzed by two-sided unpaired *t*-test or Mann-Whitney test. Multiple comparisons were analyzed by one-way ANOVA or Kruskal-Wallis tests. For survival analyses, Mantel-Cox log-rank test was performed. Significant differences are displayed as * for P ≤ 0.05, ** for P ≤ 0.01, *** for P ≤ 0.001 and **** for P ≤ 0.0001.

### Quantification of Proliferation, Cell Size and Fluorophore Intensity Measurements

Quantification of mitotic events, as measured by pHH3^+^ cells per gut, was performed by acquiring 4 independent images at 40X magnification per gut in the P3-P4 region. Images were taken in directly adjacent fields of view on each side of the gut.

#### Set 1

Quantification of mitotic events, as measured by pHH3^+^ cells per gut, was performed by acquiring 4 independent images at 40X magnification per gut in the P3-P4 region. Images were taken in directly adjacent fields of view on each side of the gut. Quantification of progenitor cell number and epithelial renewal and fluorescence intensity measurements were performed as described previously (Zipper *et al*., 2020). Fiji (ImageJ 1.51 n, Wayne Rasband, National Institutes of Health, USA) was used to calculate maximum intensity images from z-stack images. GFP positive progenitor cells and RFP positive differentiated cells (EC and/or EE) of *esg*^*ReDDM*^ (Antonello *et al*, 2015), *klu*^*ReDDM*^, and *Myo-Gal4*^*ts*^ were counted manually. Nuclear size measurements were performed in Fiji by outlining the nucleus manually and measuring the area. Mean intensities of the manually selected nuclear area were measured using Fiji by manually selecting the area of interest.

#### Set 2

Quantification of antibody fluorescence at the tSJ, bSJ and cytoplasm was performed with the ZEN Blue software. Images were acquired on a Zeiss LSM880 inverted confocal microscope equipped with AiryScan at 100X. Full z-stacks were used, and anti-Armadillo staining was used to mark membranes. 3 cells were measured per PMG. Each clearly visible tSJ in the cell was measured. 6 measurements were taken of the bicellular junction and cytoplasm in each cell, respectively.

### Smurf assay

Flies were maintained on standard medium prepared with FD&C Blue Dye no. 1 from Spectrum, added at a concentration of 2.5% (wt/vol). Loss of intestinal barrier function was determined (“Smurf” status) when dye was clearly observed outside the digestive tract (Rera *et al*., 2011). Flies were checked three times weekly for loss of barrier function and/or death.

### Survival Analysis

Flies were kept at 18°C and 3-5 days old flies were then shifted to 29°C. Not more than 10 flies (1:1 ratio of females: males) were kept per vial and flipped every other day to avoid bacterial contamination in the food. Flies were checked daily to record death events. At least 80 flies were assessed per genotype. Survival curves were plotted using Prism GraphPad software.

## Acknowledgments

The authors thank the Vienna Drosophila RNAi Center (VDRC), the Bloomington Drosophila Stock Center (NIHP400D018537), the Transgenic RNAi Project (TRiP) at Harvard Medical School (NIK/NIGMS R01-GM084947) and Thomas Klein for reagents. We also thank the UCLA MCDB/BSCRC Microscopy Core and Center for Advanced Imaging (CAi) at Heinrich Heine University (DFG INST 208/539-1 FUGG) for imaging training and facilities. In addition, we are grateful to Jones laboratory members for comments and feedback on experiments and the manuscript. Special thanks are also given to Volker Hartenstein for sharing laboratory space and equipment. This work was supported by the Eli and Edythe Broad Center of Regenerative Medicine and Stem Cell Research at the University of California, Los Angeles and the NIH: R01AG028092, R01DK105442, R01GM135767 (D.L.J.). MG is funded by the Deutsche Forschungsgemeinschaft (DFG-Sachbeihilfe RE 3453/6–1).

## Supplemental Information

**Fig EV1:**
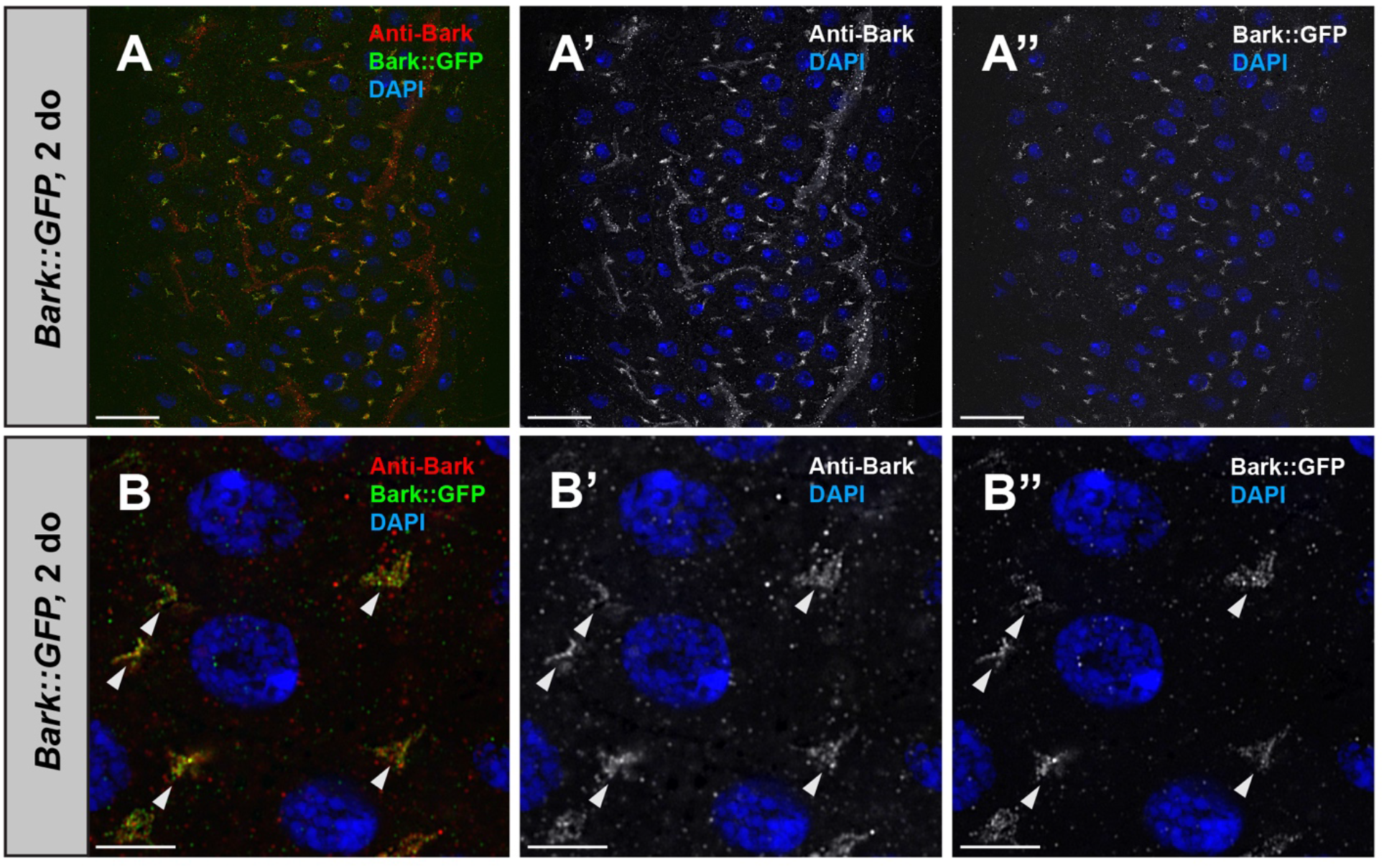
Bark is expressed at the tSJ in the adult *Drosophila* hindgut. (**A-A”**) Representative staining for Bark (antisera, red) (*Bark::GFP*, GFP, green) in the hindgut of young (2 do) flies. DNA is stained with DAPI (blue). Scale bars, 20 μm. (**B-B”**) High magnification view of the hindgut showing staining of Bark at the tSJ (arrowheads). Scale bars, 5 μm.

**Fig EV2:**
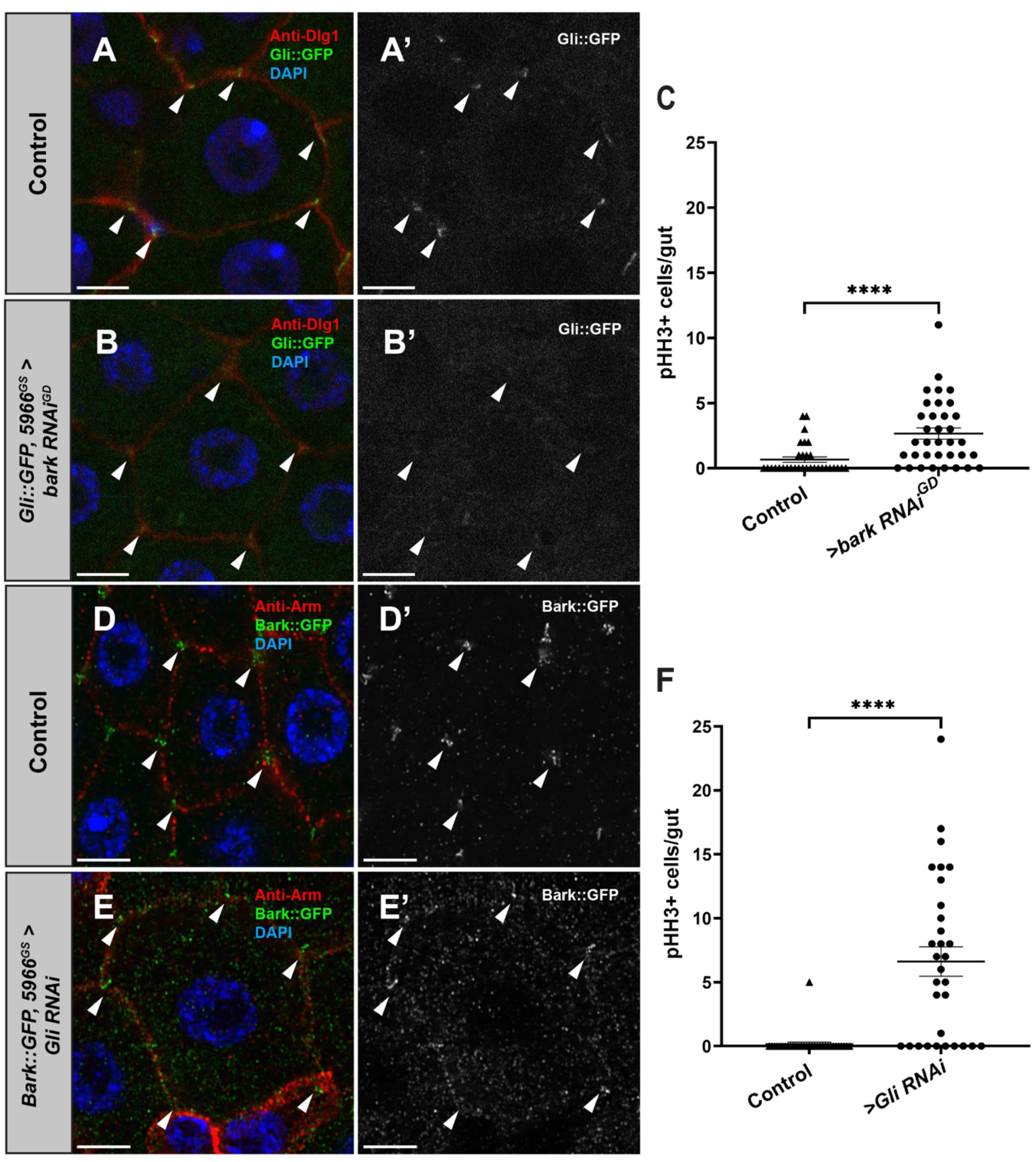
Depletion of Gli results in loss of Bark from the TCJ in the adult posterior midgut. (**A-A’**) In young outcrossed (7 do) fly PMGs, Bark is localized exclusively to the tSJ. (**B-B’**) Depletion of Gli expression in ECs of young (7 do) flies leads to loss of Bark from the tSJ and an increase in Bark on the bicellular junction and cytoplasm. Genotypes: *w; 5966*^*GS*^ (control); *w; Bark::GFP, 5966*^*GS*^*:Gli RNAi*. All flies are fed RU-486 for 5 days to induce transgene expression. Bark (GFP, green); Armadillo (Arm, adherens junctions, red); DNA (DAPI, blue). Scale bars, 20 μm. (**C**) Quantification of mitotic events in the posterior midguts of *5966*^*GS*^*:Gli RNAi* and outcrossed controls. Statistical significance was determined by Mann-Whitney test. **, *P* < .01. (**D-D’**) In young outcrossed fly PMGs, Gli localizes exclusively to the tSJ. (**E-E’**) EC-specific depletion of Bark protein by *5966*^*GS*^*:bark RNAi*^*GD*^ expression causes mislocalization of Gli to cytoplasmic puncta, indicating Bark is required for Gli localization to the TCJ in the PMG. Genotypes: *w; Gli-GFP; bark RNAi*^*GD*^ (control); *w; Gli-GFP; 5966*^*GS*^*:bark RNAi*^*GD*^. All flies are fed RU-486 for 7 days to induce transgene expression. Scale bars, 20 μm. (**F**) Quantification of mitotic events in the posterior midguts of *5966*^*GS*^*:bark RNAi*^*GD*^ and outcrossed controls. Statistical significance was determined by Mann-Whitney test. ****, *P* < .0001.

**Fig EV3:**
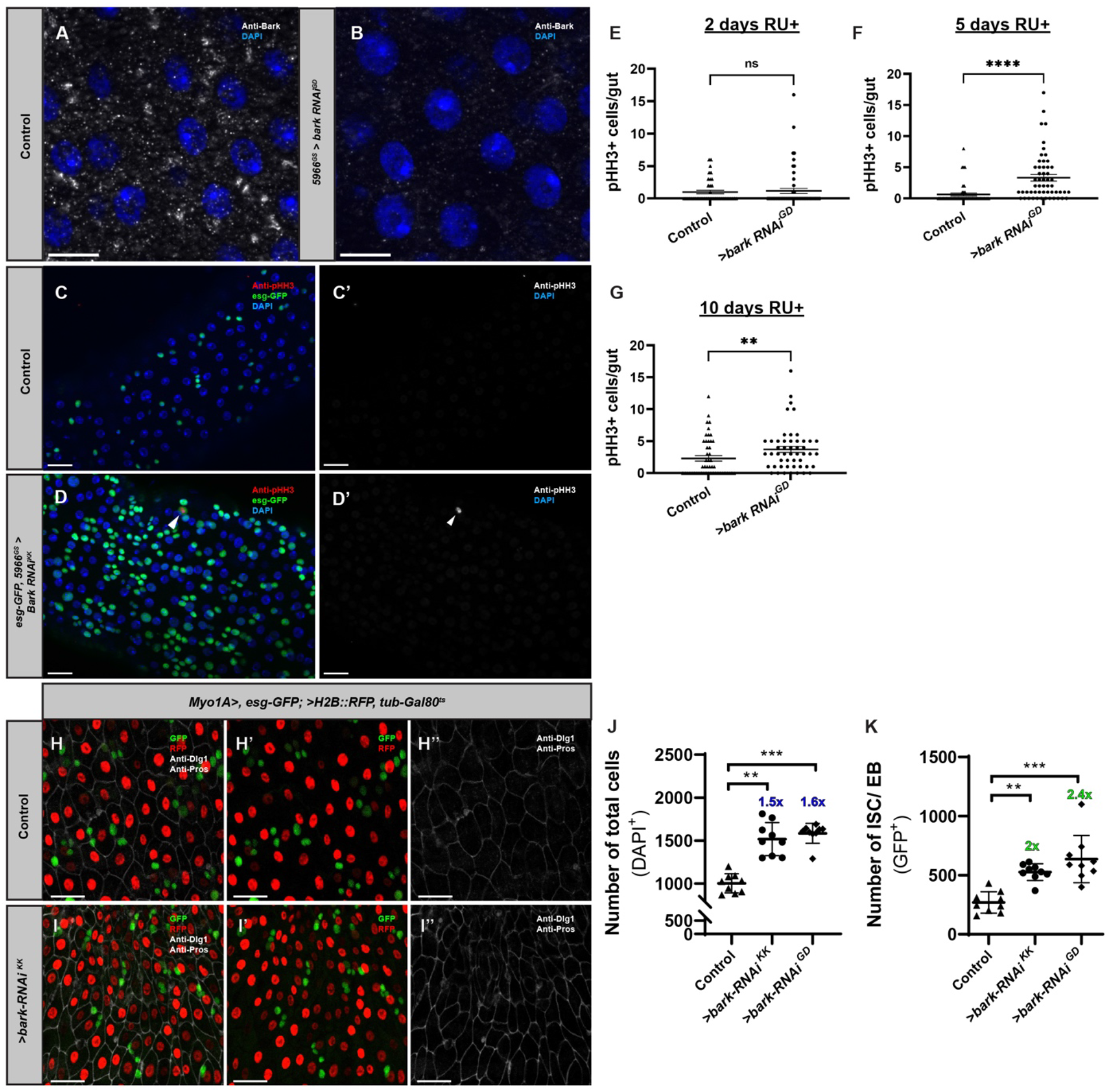
Depletion of bark in mature EC non-autonomously stimulates stem cell proliferation. Induced expression of *5966*^*GS*^*:bark RNAi*^*GD*^ by RU-486 feeding (11d) successfully depletes Bark protein from ECs in PMGs from young (2 do) flies (**B**) compared to uninduced siblings (**A**). Scale bar, 10 μm. (**C-D’**) Depletion of *bark* by induced expression of *5966*^*GS*^*:bark RNAi*^*KK*^ in young flies results in an increase in esg+ cells and mitotic events in the PMG (n = 22) compared to outcrossed controls (n = 26), similarly to *5966*^*GS*^:*bark RNAi*^*GD*^ as measured by phospho-histone H3 (arrowhead). Genotypes: *w; esg-GAL4, UAS-GFP, 5966*^*GS*^ (outcrossed control); *w; esg-GAL4, UAS-GFP, 5966*^*GS*^*:bark RNAi*^*KK*^ All flies are fed RU-486 for 5 days to induce transgene expression. Scale bars, 20 μm. (**E-G**) Depletion of Bark leads to a significant increase in ISC proliferation over time. Depletion of Bark protein from young fly PMG enterocytes by induced expression of *5966*^*GS*^*:bark RNAi*^*GD*^ results in an increase in esg+ cells and mitotic events in the PMG compared to outcrossed controls 2d (**E**), 5d (**F**) and 10d (**G**) following RU-486 induction. Genotypes: *w; esg-GAL4, UAS-GFP, 5966*^*GS*^ (outcrossed control); *w; esg-GAL4, UAS-GFP, 5966*^*GS*^*:bark RNAi*^*GD*^. 2d induction (**E**): >*bark RNAi*^*GD*^ n = 58; *w*^*1118*^ n = 36. 5d induction (**F**): >*bark RNAi*^*GD*^ n = 54; *w*^*1118*^ n = 48. 10d induction (**G**): >*bark RNAi*^*GD*^ n = 50; *w*^*1118*^ n = 49. Data shown is combined from two independent trials. Error bars represent mean with SEM. Significance was determined by Mann-Whitney test. ns, not significant; **, *P* < .005; ****, *P* < .0001. Knockdown of *bark* in ECs using *Myo1A-GAL4*^*ts*^ (*Myo1A>, esg-GFP; H2B::RFP, tub-GAL80*^*ts*^) increases progenitor cells. **(H-H’’)** Confocal images of control PMG in the R5 region after 7 days of tracing. **(I-I’’)** Knockdown of *bark* increases the total number of cells and the number of ISC/ EB (GFP^+^). Anti-Dlg1 marks bSJs and anti-Pros marks EE nuclei. **(J, K)** Quantifications of the total cell number (DAPI^+^) **(J)** and ISC/ EB **(K)** upon *bark* knockdown after 7 days of tracing. (control n = 9, *>bark RNAi*^*KK*^ n = 9, *>bark RNAi*^*GD*^ n = 9). Statistical significance determined by Kruskal-Wallis test. **, *P* < .01; ***, *P* < .001. Scale bar, 20 μm.

**Fig EV4:**
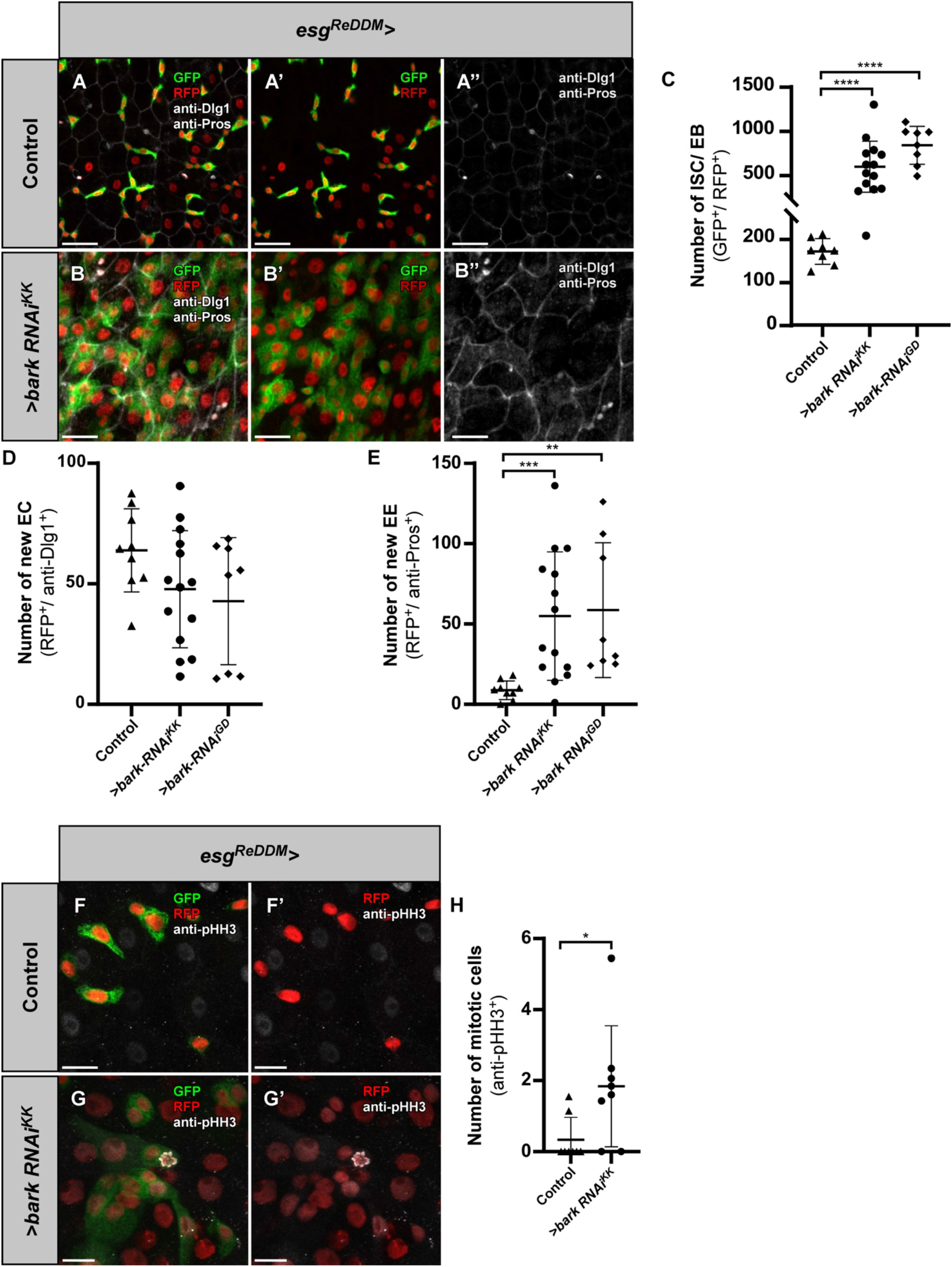
ISC/EB specific depletion of *bark* results in ISC proliferation and increase in progenitor cells and EE differentiation. *esg*^*ReDDM*^ (*w; esg-GAL4, UAS-CD8::GFP; UAS-H2B::RFP, tub-GAL80*^*ts*^) tracing and the knockdown of *bark* in ISCs/EBs leads to an increase in progenitor cells and EEs. **(B-B’’)** Compared with controls **(A-A”)**, knockdown of *bark* in the R5 region following 7d of temperature shift increases the number of newly differentiated EEs (GFP^-^/RFP^+^/anti-Pros^+^) and the number of progenitor cells (GFP^+^/RFP^+^) but no significant change in the number of ECs (GFP^-^/RFP^+^/anti-Dlg1^+^). **(C-E)** Quantifications of ISCs/EBs **(C)**, new ECs **(D)** and new EEs. **(E)** Quantification of new ECs upon *bark* knockdown after 7 days of tracing. (control n = 9, *>bark RNAi*^*KK*^ n = 14, *>bark RNAi*^*GD*^ n = 8). Statistical significance determined by one-way ANOVA. **, *P* < .01; ***, *P* < .001. Scale bar, 20 μm. **(F-G’)** Knockdown of *bark* **(G-G’)** leads to an increase in the number of proliferating cells (marked by pHH3) as compared to the controls **(F-F’). (H)** Quantification of the number of mitotic cells upon *bark* depletion after 7 days of tracing. (control n = 8, *>bark RNAi*^*KK*^ n = 8). Statistical significance determined by Mann-Whitney test. ****, *P* < .0001. Scale bar 10 μm.

## References

Antonello Z.A., Reiff T., Ballesta-Illan E. & Dominguez M. (2015) Robust intestinal homeostasis relies on cellular plasticity in enteroblasts mediated by miR-8–Escargot switch. EMBO J 34: 2025–2041

Apidianakis Y. & Rahme L.G. (2011) Drosophila melanogaster as a model for human intestinal infection and pathology. Dis Model Mech 4: 21

Biteau B., Hochmuth C.E. & Jasper H. (2008) JNK Activity in Somatic Stem Cells Causes Loss of Tissue Homeostasis in the Aging Drosophila Gut. Cell Stem Cell 3: 442–455

Biteau B., Karpac J., Supoyo S., DeGennaro M., Lehmann R. & Jasper H. (2010) Lifespan Extension by Preserving Proliferative Homeostasis in Drosophila. PLoS Genet 6: e1001159

Brand A.H. & Perrimon N. (1993) Targeted gene expression as a means of altering cell fates and generating dominant phenotypes. Development 118: 401–415

Byri S., Misra T., Syed Z.A., Bätz T., Shah J., Boril L., Glashauser J., Aegerter-Wilmsen T., Matzat T., Moussian B., et al. (2015) The Triple-Repeat Protein Anakonda Controls Epithelial Tricellular Junction Formation in Drosophila. Dev Cell 33: 535–548

Chen J., Sayadian A., Lowe N., Lovegrove H.E. & St. Johnston D. (2018) An alternative mode of epithelial polarity in the Drosophila midgut. PLoS Biol 16: e3000041

Chen J. & St. Johnston D. (2022) De novo apical domain formation inside the Drosophila adult midgut epithelium. eLife 11:e76366. DOI: https://doi.org/10.7554/eLife.76366.

Clark R.I., Salazar A., Yamada R., Fitz-Gibbon S., Morselli M., Alcaraz J., Rana A., Rera M., Pellegrini M., Ja W.W., et al. (2015) Distinct Shifts in Microbiota Composition during Drosophila Aging Impair Intestinal Function and Drive Mortality. Cell Rep 12: 1656–67

Dutta D., Dobson A.J., Houtz P.L., Gläßer C., Revah J., Korzelius J., Patel P.H., Edgar B.A. & Buchon N. (2015) Regional Cell-Specific Transcriptome Mapping Reveals Regulatory Complexity in the Adult Drosophila Midgut. Cell Rep 12: 346–358

Esmangart de Bournonville T. & le Borgne R. (2020) Interplay between Anakonda, Gliotactin, and M6 for Tricellular Junction Assembly and Anchoring of Septate Junctions in Drosophila Epithelium. Current Biology 30: 4245-4253.e4

Furriols M. & Bray S. (2000) Dissecting the Mechanisms of Suppressor of Hairless Function. Dev Biol 227: 520–532

Furuse M. & Tsukita S. (2006) Claudins in occluding junctions of humans and flies. Trends Cell Biol 16: 181–188

Guo L., Karpac J., Tran S.L. & Jasper H. (2014) PGRP-SC2 Promotes Gut Immune Homeostasis to Limit Commensal Dysbiosis and Extend Lifespan. Cell 156: 109–122

Hildebrandt A., Pflanz R., Behr M., Tarp T., Riedel D. & Schuh R. (2015) Bark beetle controls epithelial morphogenesis by septate junction maturation in Drosophila. Dev Biol 400: 237–247

Hung R.J., Li J.S.S., Liu Y. & Perrimon N. (2021) Defining cell types and lineage in the Drosophila midgut using single cell transcriptomics. Curr Opin Insect Sci 47: 12–17

Izumi Y., Furuse K. & Furuse M. (2019) Septate junctions regulate gut homeostasis through regulation of stem cell proliferation and enterocyte behavior in Drosophila. J Cell Sci: jcs.232108

Izumi Y. & Furuse M. (2014) Molecular organization and function of invertebrate occluding junctions. Semin Cell Dev Biol 36: 186–193

Jiang H., Patel P.H., Kohlmaier A., Grenley M.O., McEwen D.G. & Edgar B.A. (2009) Cytokine/Jak/Stat signaling mediates regeneration and homeostasis in the Drosophila midgut. Cell 137: 1343

Kirkwood T.B.L. (2004) Intrinsic ageing of gut epithelial stem cells. Mech Ageing Dev 125: 911–915

Korzelius J., Azami S., Ronnen-Oron T., Koch P., Baldauf M., Meier E., Rodriguez-Fernandez I.A., Groth M., Sousa-Victor P. & Jasper H. (2019) The WT1-like transcription factor Klumpfuss maintains lineage commitment of enterocyte progenitors in the Drosophila intestine. Nature Communications 2019 10:1 10: 1–13

Lane N.J. & Skaer H. (1980) Intercellular Junctions in Insect Tissues. In pp 35–213.

Li H. & Jasper H. (2016) Gastrointestinal stem cells in health and disease: from flies to humans. Dis Model Mech 9: 487–99

Marchiando A.M., Graham W.V. & Turner J.R. (2010) Epithelial Barriers in Homeostasis and Disease. Annual Review of Pathology: Mechanisms of Disease 5: 119–144

Mariano C., Sasaki H., Brites D. & Brito M.A. (2011) A look at tricellulin and its role in tight junction formation and maintenance. Eur J Cell Biol 90: 787–796

Micchelli C.A. & Perrimon N. (2006) Evidence that stem cells reside in the adult Drosophila midgut epithelium. Nature 439: 475–479

Ohlstein B. & Spradling A. (2005) The adult Drosophila posterior midgut is maintained by pluripotent stem cells. Nature 2005 439:7075 439: 470–474

Ohlstein B. & Spradling A. (2007) Multipotent Drosophila intestinal stem cells specify daughter cell fates by differential Notch signaling. Science 315: 988–992

Osterwalder T., Yoon K.S., White B.H. & Keshishian H. (2001) A conditional tissue-specific transgene expression system using inducible GAL4. Proc Natl Acad Sci U S A 98: 12596–601

Park J., Kim Y. & Yoo M. (2009) The role of p38b MAPK in age-related modulation of intestinal stem cell proliferation and differentiation in Drosophila. Aging 1: 637

Reiff T., Antonello Z.A., Ballesta-Illán E., Mira L., Sala S., Navarro M., Martinez L.M. & Dominguez M. (2019) Notch and EGFR regulate apoptosis in progenitor cells to ensure gut homeostasis in Drosophila. EMBO J 38: e101346

Ren W., Wu K., Li X., Luo M., Liu H., Zhang S. & Hu Y. (2014) Age-related changes in small intestinal mucosa epithelium architecture and epithelial tight junction in rat models. Aging Clin Exp Res 26: 183–191

Rera M., Bahadorani S., Cho J., Koehler C.L., Ulgherait M., Hur J.H., Ansari W.S., Lo T., Jones D.L. & Walker D.W. (2011) Modulation of longevity and tissue homeostasis by the Drosophila PGC-1 homolog. Cell Metab 14: 623–634

Rera M., Clark R.I. & Walker D.W. (2012) Intestinal barrier dysfunction links metabolic and inflammatory markers of aging to death in Drosophila. Proc Natl Acad Sci U S A 109: 21528–33

Resnik-Docampo M., Koehler C.L., Clark R.I., Schinaman J.M., Sauer V., Wong D.M., Lewis S., D’Alterio C., Walker D.W. & Jones D.L. (2017) Tricellular junctions regulate intestinal stem cell behaviour to maintain homeostasis. Nat Cell Biol 19: 52–59

Resnik-Docampo M., Sauer V., Schinaman J.M., Clark R.I., Walker D.W. & Jones D.L. (2018) Keeping it tight: The relationship between bacterial dysbiosis, septate junctions, and the intestinal barrier in Drosophila. Fly (Austin) 12: 34–40

Salazar A.M., Resnik-Docampo M., Ulgherait M., Clark R.I., Shirasu-Hiza M., Jones D.L. & Walker D.W. (2018) Intestinal Snakeskin Limits Microbial Dysbiosis during Aging and Promotes Longevity. iScience 9: 229–243

Sarov M., Barz C., Jambor H., Hein M.Y., Schmied C., Suchold D., Stender B., Janosch S., Vinay Vikas K.J., Krishnan R.T., et al. (2016) A genome-wide resource for the analysis of protein localisation in Drosophila. Elife 5

Schiffrin E.J., Morley J.E., Donnet-Hughes A. & Guigoz Y. (2010) The inflammatory status of the elderly: The intestinal contribution. Mutation Research/Fundamental and Molecular Mechanisms of Mutagenesis 690: 50–56

Tepass U. & Hartenstein V. (1994) The Development of Cellular Junctions in the Drosophila Embryo. Dev Biol 161: 563–596

Tran L. & Greenwood-Van Meerveld B. (2013) Age-Associated Remodeling of the Intestinal Epithelial Barrier. J Gerontol A Biol Sci Med Sci 68: 1045

Wittek A., Hollmann M., Schleutker R. & Luschnig S. (2020) The Transmembrane Proteins M6 and Anakonda Cooperate to Initiate Tricellular Junction Assembly in Epithelia of Drosophila. Current Biology 30: 4254-4262.e5

Choi Y.J., Hwang M.S., Park J.S., Bae S.K., Kim Y.S., Yoo M.A. (2008) Age-related upregulation of Drosophila caudal gene via NF-kappaB in the adult posterior midgut. Biochim Biophys Acta 1780: 1093–1100

Zhai Z., Kondo S., Ha N., Boquete J.P., Brunner M., Ueda R. & Lemaitre B. (2015) Accumulation of differentiating intestinal stem cell progenies drives tumorigenesis. Nature Communications 2015 6:1 6: 1–13

Zipper L, Jassmann D, Burgmer S, Görlich B & Reiff T (2020) Ecdysone steroid hormone remote controls intestinal stem cell fate decisions via the PPARγ-homolog Eip75B in Drosophila. Elife 9: 1–27

